# A *cis*-regulatory point mutation at a R2R3-Myb transcription factor contributes to speciation by reinforcement in *Phlox drummondii*

**DOI:** 10.1101/2023.04.19.537550

**Authors:** Austin G. Garner, Andrew Cameron, Andrea E. Berardi, Robin Hopkins

## Abstract

The process of reinforcement, whereby selection favors the evolution of increased reproductive trait divergence to reduce costly hybridization between species, has been well documented in nature, yet we know very little about how this process evolves at the molecular level. In this study, we combine functional characterization and genetic association tests to identify the mutational basis of reinforcement in the Texas wildflower *Phlox drummondii. P. drummondii* evolved from light to dark flower color intensity by selection to stop hybridization with the closely related species *P. cuspidata*, and previous research suggests differential expression of a R2R3-Myb transcription factor underlies this phenotypic transition. Using gene-silencing experiments, we demonstrate expression of this transcription factor does control variation in flower color intensity. We then apply association mapping across a large genomic region flanking the R2R3-Myb gene and identified a point mutation within the gene’s promoter that is highly associated with flower color intensity in nature. Alleles at this mutation site match the expected patterns of dominance, create variation in predicted cis-regulatory motifs within the R2R3-Myb proximal promoter, and occur in the direction of evolution predicted for flower color variation in this system. By identifying the mutational basis of reinforcement in this system we demonstrate that, as predicted by theory, reproductive isolation can evolve despite gene flow through a very simple genetic basis.

## Introduction

Natural selection can facilitate speciation through the process of reinforcement, whereby an increase in reproductive isolation between diverging lineages is favored by selection to decrease costly hybridization (Dobzhansky, 1937). Evidence of reinforcement in the process of speciation has been documented throughout the tree of life. (Butlin, 1987; Howard, 1993; Servedio & Noor, 2003; Ortiz-Barrientos *et al*., 2009; Garner *et al*., 2018). However, how reinforcement evolves at the molecular level remains largely unknown. The formalization of population genetics during the Modern Synthesis led to the idea that identifying the genetic basis of barriers to reproduction could better our understanding of how these barriers evolve and the process of speciation; thus far this notion has proven true (Dobzhansky, 1936, 1937; Coyne & Orr, 2004; Bomblies & Peichel, 2022). Much less is known about the genetic basis of pre-zygotic reproductive isolation, and specifically reinforcement, than about post-zygotic barriers to reproduction. By functionally demonstrating the genes causing reinforcement and isolating the mutations underlying their natural variation, we can address long-standing questions about how reinforcement occurs in nature.

Due to the historical controversy about the feasibility of reinforcement, extensive theoretical research has characterized the evolutionary conditions under which reinforcement is likely to occur. (Kirkpatrick & Ravigné, 2002; Servedio & Noor, 2003). A primary argument against the feasibility of reinforcement is that the hybridization driving reinforcing selection can also lead to species fusion, if gene flow is too extensive, or extinction, if hybridization is too costly (Felsenstein, 1981). Reinforcement can only successfully evolve when the mutations contributing to increased reproductive isolation can rapidly rise in frequency within a species without becoming disassociated from species-specific adaptations or genetic backgrounds by interspecific gene flow and recombination (Servedio & Noor, 2003; Coyne & Orr, 2004). Mathematical theory proposes, one solution to this problem is for reinforcement to be caused by few loci with large phenotypic effects, such that selection is strong enough on each locus to allow for evolution within a lineage without disassociation (Felsenstein, 1981; Kirkpatrick, 2000). However, we have little to no empirical data to evaluate this expectation. This solution could be accomplished by a single large effect mutation or be the combination of multiple interacting polymorphisms with varied effect sizes in linkage (Kirkpatrick & Barton, 2006; Yeaman & Whitlock, 2011; Remington, 2015). Identifying the contributions of individual genes and mutations to traits that evolved by reinforcement is necessary to understand which of these theoretical genetic mechanisms has favored the evolution of reinforcement in nature. Additionally, this knowledge will inform if reinforcement evolves by similar or different molecular pathways and constraints as selection for other adaptive traits.

Variation in *Phlox drummondii* flower color is one of the best studied cases of reinforcement to date. *P. drummondii* and the closely related species *P. cuspidata* both display similar light-blue colored flowers where their natural ranges do not overlap (allopatry); however, where these two species co-occur (sympatry), *P. drummondii* has evolved dark-red flower color (Levin, 1985). *P. drummondii and P. cuspidata* can be frequently found together in sympatry and both species share visitation by a common pollinator, the *Battus philenor* butterfly (Hopkins & Rausher, 2012; Briggs *et al*., 2018). Pollen exchange between these two species is costly due to the formation of nearly sterile hybrid offspring, and genomic evidence confirms there is a history of hybridization and gene flow where they coexist. (Levin, 1985; Ferguson *et al*., 1999; Roda *et al*., 2017; Suni & Hopkins, 2018; Garner *et al*., 2022). Our previous work demonstrates that the evolution of dark *P. drummondii* flower color in sympatry evolved under selection by reinforcement to prevent hybridization with *P. cuspidata* (Hopkins & Rausher, 2012; Hopkins *et al*., 2014). *P. drummondii* with dark flower color induces floral constancy in the foraging behavior of the *B. philenor* butterfly, reducing the rate of pollen exchange and costly hybridization between *P. cuspidata* and *P. drummondii* by 50%, relative to when *P. drummondii* has light flower color (Hopkins & Rausher, 2012). Associations from controlled crosses suggest that the large phenotypic transition in *P. drummondii* flower intensity (from light to dark) may be caused by unidentified dominant *cis*-regulatory mutation(s) at a R2R3-Myb gene, a class of transcription factor that can mediate production of anthocyanin pigmentation in flowers (Hopkins & Rausher, 2011). Here we refer to this gene as *PdMyb1*.

Given our understanding of the role of flower intensity in the reinforcement of *P. drummondii* and its likely genetic basis, *P. drummondii* is a prime system for dissecting the mutational basis of reinforcement so we can better understand how this process evolves. In this study, we leverage natural genetic variation in *P. drummondii* flower color to functionally demonstrate the contribution of *PdMyb1* in the divergence from light to dark flower color and to infer the mutational and evolutionary history by which dark flower color evolved by reinforcement.

### Natural variation in *P. drummondii* flower color

We first evaluated if the natural variation in flower color found throughout the range of *P. drummondii* can best be summarized by four-color categories previously identified using controlled genetic crosses. Previous crosses between the two predominant flower color phenotypes in *P. drummondii* (light-blue and dark-read) reveal that flower color variation can be caused by two loci with two alleles each, resulting in four discrete flower color categories (i.e., light-red, light-blue, dark-red, and dark-blue) (Figure 1a) (Hopkins & Rausher, 2011). One locus corresponds to variation in flower color hue, either blue or red, and determines the type of floral pigments produced, and the second locus corresponds to variation in flower color intensity, either light or dark, and determines the amount of pigment produced in the flower. Genotype and expression of *PdMyb1* is associated with the transition from light to dark flower color in this cross (Hopkins & Rausher, 2011). Light-blue allopatric and dark-red sympatric *P. drummondii* are fully interfertile and generate natural admixed zones in a narrow region of contact between sympatry and allopatry (Hopkins *et al*., 2014) (Figure 1b). We leveraged the natural segregation of flower color within and around these contact zones to describe the flower color variation in *P. drummondii* and determine if a two-locus model is adequate to explain the genetic variation in flower color observed throughout the species’ range.

**Figure 1:**
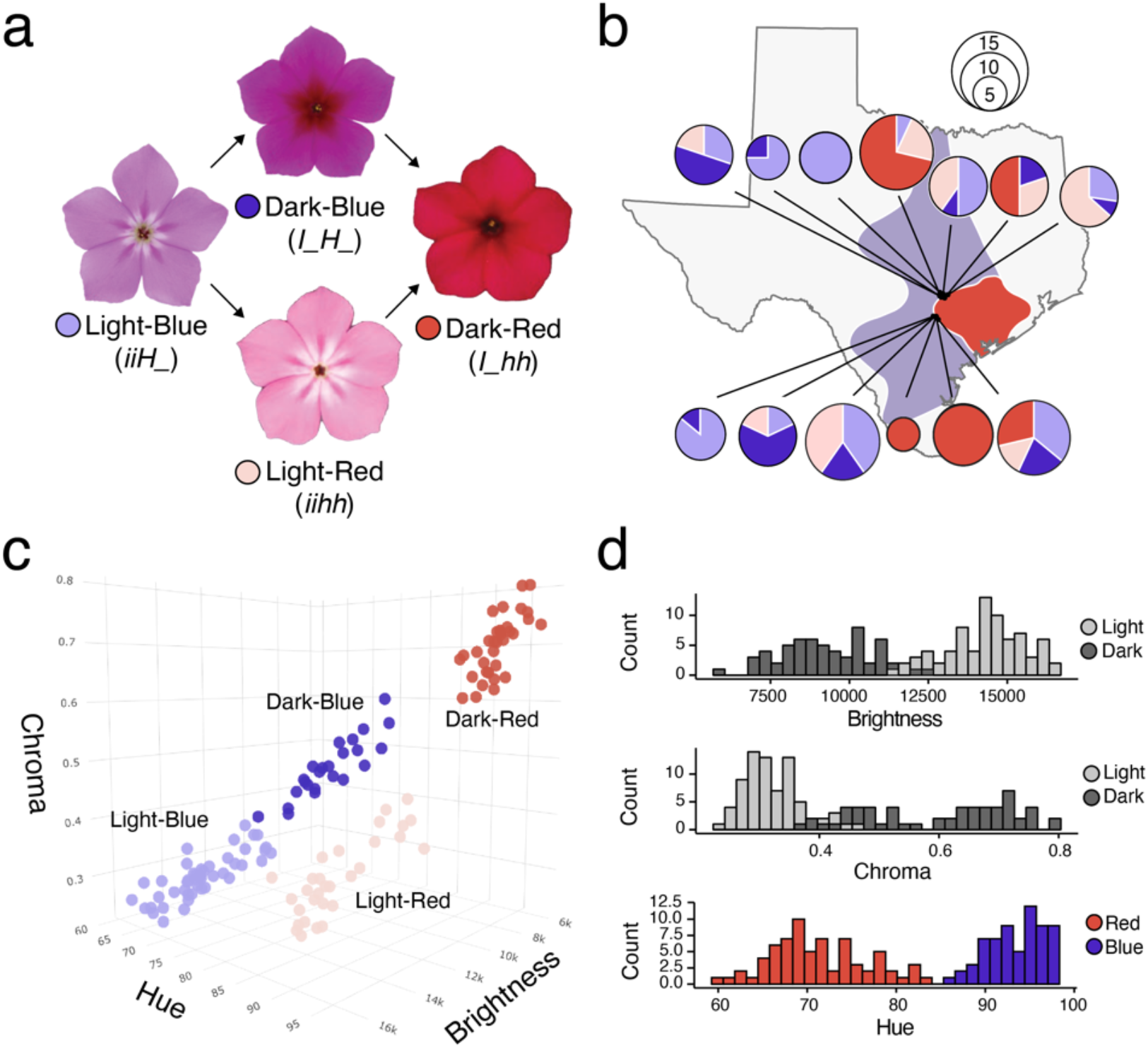
Two large evolutionary transitions explain flower color variation in natural admixed *P. drummondii* populations. **a**. Representative images of the four flower color categories previously inferred in controlled crosses suggesting a large evolutionary transition in intensity (light to dark) and hue (blue to red) in the reinforcement of dark-red flower color. Predicted hue (H/h) and intensity (I/i) genotypes are indicated below each flower color category, with dominant alleles capitalized. **b**. Schematic of Texas showing the sampling locations of natural admixed individuals from two independent zones of sympatry-allopatry contact included in this study. Pie charts show the number of individuals of each color category collected from each sampling site. The underlying light-blue and dark-red polygons indicate the allopatric and sympatric *P. drummondii* distributions, respectively. **c**. Coordinates of sampled admixed individuals along three-dimensions of spectral flower color (brightness, chroma, and hue). Color indicates individual assignment to one of the four model-inferred color categories (light-blue, dark-blue, light-red, and dark-red). **d**. Histograms of the values of brightness, chroma, and hue in the admixed individuals, with individual color assignment based on the categorical variable, intensity (light or dark) or hue (red or blue), that best explains variation along each color axis.

We characterized the flower color of 134 admixed individuals, from two independent sympatry-allopatry contact zones, both quantitatively using floral spectral reflectance and qualitatively by eye, into one of the four flower color categories (Figure 1b). We decomposed the spectral reflectance measurements into three quantitative axes of color: hue, brightness, and chroma. Analysis of these color values with unsupervised gaussian mixture modeling determined the natural flower color variation in these admixed individuals is best described as a mixture of four gaussian components or “clusters” (Figure 1c, Supplemental Information Figure S1, S2). The means and standard errors of the color axes in this four-component model indicate the presence of a light (high brightness, low chroma) and a dark (low brightness, high chroma) component in both the red and blue ranges of hue (Supplemental Information

Table S3.3). Categorical classification of the model components by light or dark explains most of the natural phenotypic variance in chroma (71.3%) and brightness (78.4%), while classification by red or blue components strongly explains the natural phenotypic variance in hue (86.4%) (Figure 1d, Supplemental Information Figure 3). Comparison of the model-inferred and our qualitative light-red, dark-red, light-blue, dark-blue group membership demonstrates they are nearly identical (Adjusted Rand Index=0.98; X-squared = 393.84, df = 9, p-value < 2.2e-16), except for one individual that the model classified with low confidence. Comparison of the spectral reflectance color axis values of the ancestral (light-blue) and derived (dark-red) flower colors observed in allopatry and sympatry, respectively, show they are indistinguishable from the light-blue and dark-red phenotypes observed in the admixed zone (Supplemental Information Figure S4).

Our results demonstrate natural variation in *P. drummondii* flower color segregates into the four discrete categories previously identified in a controlled cross study and support the evolution of dark-red *P. drummondii* flower color can be classified as a discrete transition in intensity (light to dark) and hue (blue to red). Our nearly identical replication of the phenotypic partitioning observed in the earlier controlled cross study suggests natural variation in *P. drummondii* flower may be controlled by the same two large effect loci inferred in the crossing study, with intensity potentially being regulated by the associated gene *PdMyb1*.

### Differential expression of *PdMyb1* causes variation in *P. drummondii* flower intensity

*PdMyb1* gene expression has been previously shown to correlate with flower color intensity and total quantity of floral anthocyanin pigment (Hopkins & Rausher, 2011). R2R3-Myb transcription factors regulate a variety of plant biological processes, including secondary metabolism, development, and resistance to abiotic and biotic stress (Dubos *et al*., 2010). However, the Subgroup 6 R2R3-Myb family is known to regulate anthocyanin biosynthesis and can cause variation in the amount and patterning of floral anthocyanin pigmentation (Stracke *et al*., 2001; Dubos *et al*., 2010). To determine if *PdMyb1* is a Subgroup 6 R2R3-Myb, we isolated the full amino acid sequence of the *PdMyb1* gene model from the *P. drummondii* reference genome and aligned it to a taxonomically diverse set of known anthocyanin-regulating Subgroup 6 R2R3-Myb gene sequences. *PdMyb1* contains well-known conserved amino acid motifs of the Subgroup 6 R2R3-Myb gene family, supporting the hypothesis that *PdMyb1* is an anthocyanin regulating R2R3-Myb gene (Supplemental Information Figure S5).

To validate the functional role of *PdMyb1* transcript abundance in regulating flower color intensity, we used virus-induced gene silencing (VIGS) to knock down *PdMyb1* transcript levels in naturally dark flowered plants (Lu *et al*., 2003) (Figure 2a). Using quantitative real-time PCR, we verify successful silencing of *PdMyb1* on some branches of our infected plants while other branches remained unsilenced and serve here as internal controls (Figure 2b). *PdMyb1* silencing produces flowers that are significantly lighter in flower intensity (higher brightness, lower chroma) relative to unsilenced flowers from the same plant (Figure 2c-d). *PdMyb1* unsilenced flowers do not differ in *PdMyb1* expression or flower color from untreated dark plants (Supplemental Table S5c). Additionally, *PdMyb1* silenced flowers do differ in *PdMyb1* expression and flower intensity from untreated dark plants and appear more similar to *PdMyb1* expression and flower intensity of untreated light plants (Figure 2e-g).

**Figure 2:**
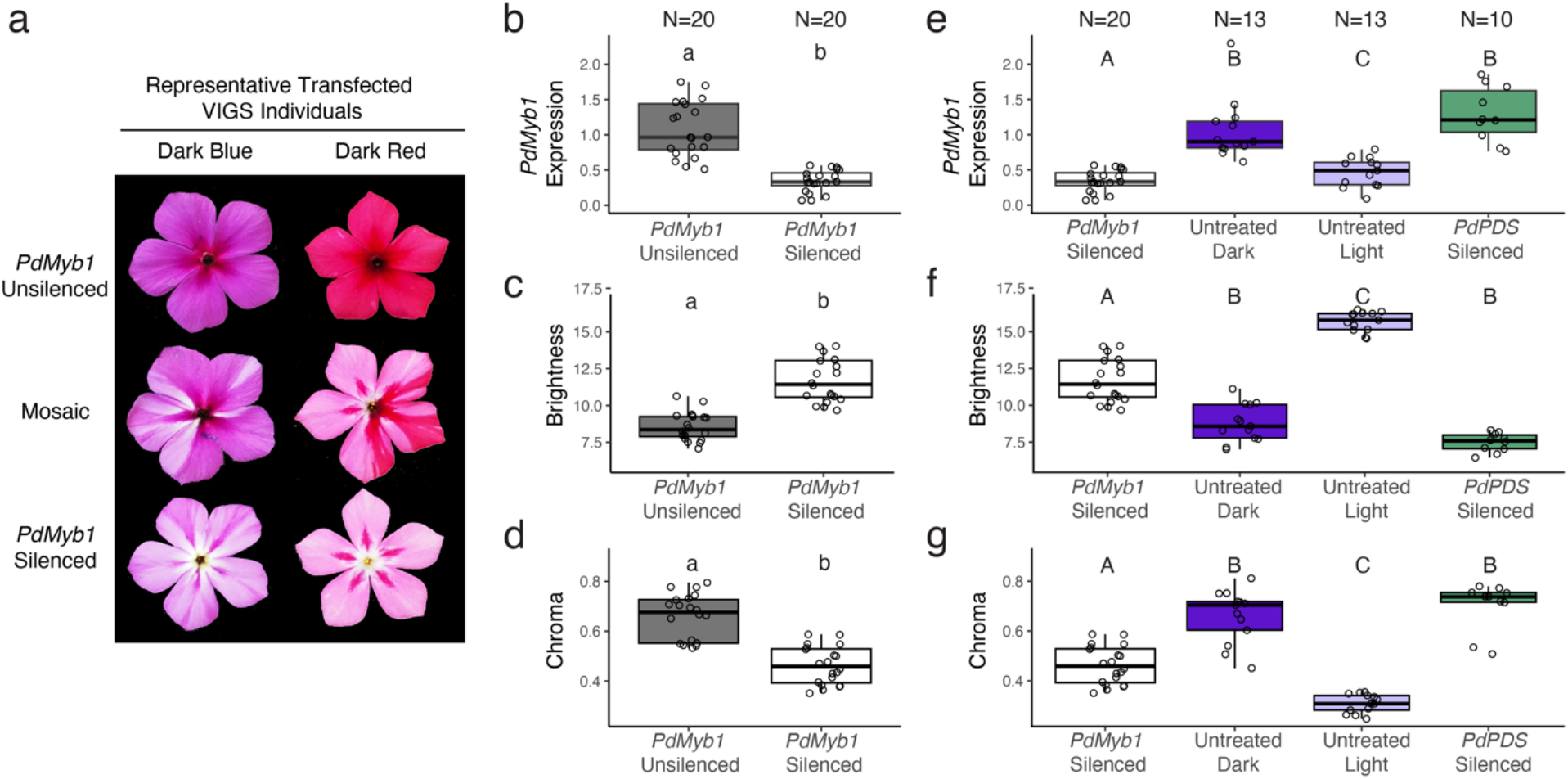
Knockdown of *PdMyb1* causes a shift in flower color intensity. **a**. Branch-specific *PdMyb1* virus-induced gene silencing (VIGS) results in *PdMyb1* unsilenced dark flowers (Top), *PdMyb1* silenced light flowers (Bottom), and flowers with mosaic *PdMyb1* silencing (Middle) on the same naturally dark flowered plants. **b-d**. *PdMyb1* VIGS silencing knocks down flower petal expression of *PdMyb1* (**b**) and creates lighter (brighter (**c**) and less chromatic (**d**)) flowers relative to *PdMyb1* unsilenced dark flowers from the same plant. VIGS-induced shifts in *PdMyb1* expression and flower intensity in dark plants is in the same direction as the divergence between natural light and dark flowered plants (**e-g**). Significant differences assessed by paired t-tests (lowercase letters) and ANOVAs with post-hoc Tukey (uppercase letters), P < 0.001 (Supplemental Information Table S3.5a-c, S3.6).

**Figure 3:**
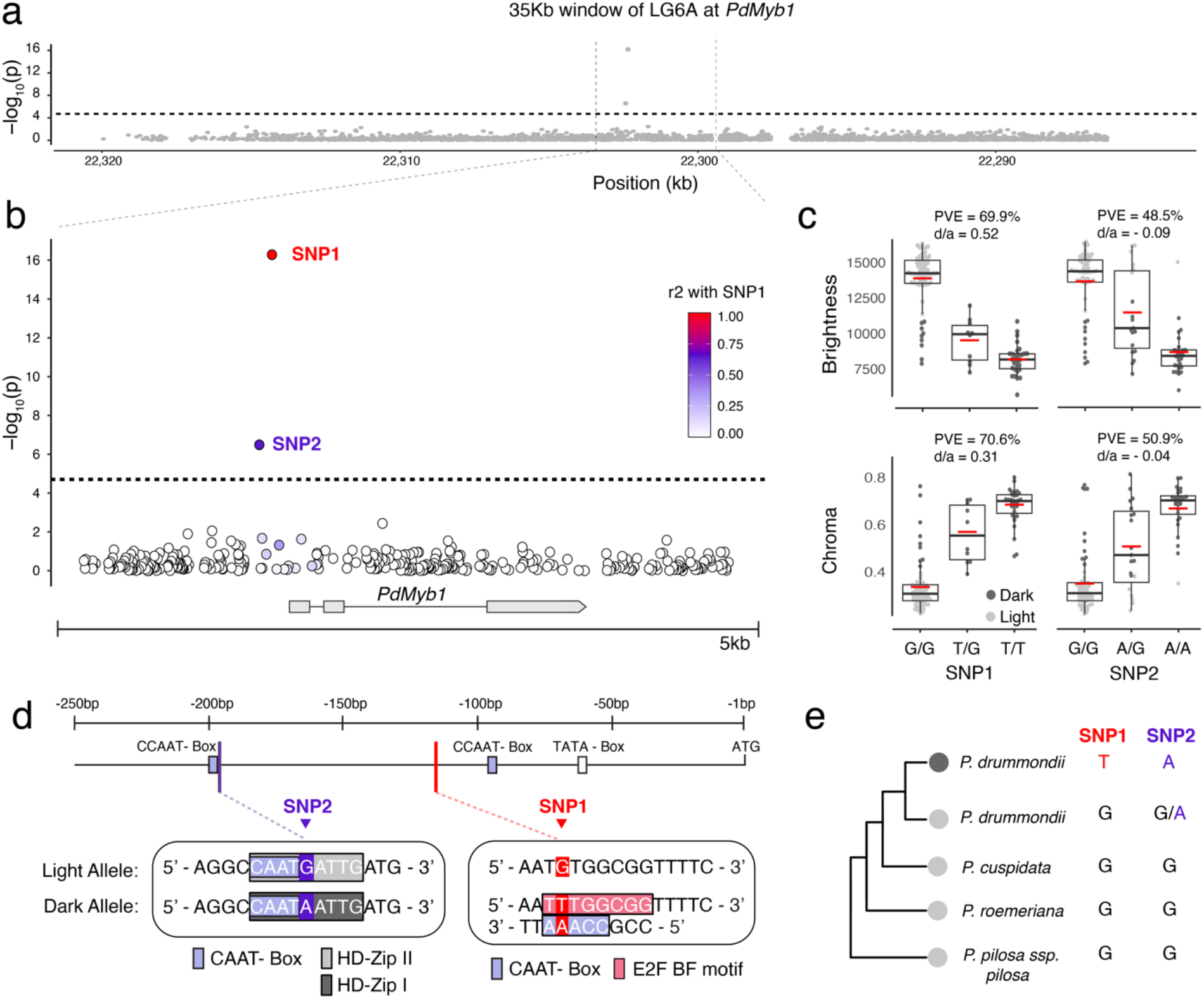
Genotype-phenotype associations and the evolution of *cis*-regulation at the *PdMyb1* locus. **a**. Targeted GWAS identifies a single peak significantly associated with natural variation in *P. drummondii* categorical flower intensity across the 35kb window flanking *PdMyb1*. The dashed line represents the Bonferroni corrected significance threshold ((-log_10_(P) ≃4.7) **b**. A closer view of two associated variants reveals they are both SNPs directly upstream of *PdMyb1* coding sequence. Circles represent genetic variants. Colors represent the degree of linkage disequilibrium between each marker and the highest associated variant, SNP1. **c**. Quantitative brightness (top) and chroma (bottom) by genotype at SNP1. Red bars denote the mean trait value for each genotype. Statistics above each plot report the proportion of variance explained (PVE) and the dominance effect (d/a) of alleles at SNP1 **d**. *Cis*-regulatory motif prediction of 250bp upstream of *PdMyb1*. SNP1 and SNP2 are located between the TATA-Box and CAAT-Box elements of the *PdMyb1* core promoter, denoted with white and light purple boxes respectively (Top). Base pair differences at SNP1 and SNP2 cause allele-specific variation at predicted regulatory motifs (Bottom). **e**. Evolutionary history at SNP1 and SNP2 confirms the dark associated alleles are derived in *P. drummondii*.

These results demonstrate that *PdMyb1* transcript abundance can cause variation in total anthocyanin production and flower color intensity along the same axes of divergence observed in natural *P. drummondii* populations. Our results are consistent with the evolutionary transition from light to dark flower intensity being caused by variation in expression of *PdMyb1*.

### Point mutations at *PdMyb1* are associated with natural variation in flower intensity

We used a region-specific association mapping approach to isolate mutations causing natural variation in flower color intensity. Previous analysis of allele-specific expression in individuals heterozygous for the light and dark flower color allele demonstrated that natural variation in *PdMyb1* expression is associated with *cis*-regulatory mutation(s) (Hopkins & Rausher, 2011). *Cis*-regulatory polymorphisms causing functional variation in the expression of R2R3-Myb transcription factors are often within 5kb of the gene (Sheehan *et al*., 2016; Li *et al*., 2022). Therefore, we hypothesized that expression differences in *PdMyb1* are due to proximal mutations. We first targeted, isolated, and long-read sequenced high molecular weight DNA corresponding to a genomic window flanking *PdMyb1* from 105 of our phenotyped wild collected individuals (82 admixed, 11 sympatric, 12 allopatric; 54 light, 51 dark). After filtering, we retained 2,533 high-quality polymorphic markers densely spanning a ∼35kb window flanking *PdMyb1*. We identify two variants associated with categorical and quantitative differences in flower color intensity that exceed the region-wide significance threshold (-log10(P-value) ≃4.7, Bonferroni corrected) (Figure 3a, Supplemental Figure S6).

Both associated variants are single nucleotide polymorphisms (SNPs) in moderately high linkage disequilibrium (LD) (r^2^=0.65) and sit 116bp and 194bp upstream of inferred the *PdMyb1* start codon (Figure 3b). Respectively, SNP1 and SNP2 explain 69.9% and 48.5% of variance in brightness and 70.6% and 50.9% of variance in chroma (Figure 3c), with SNP1 showing the strongest association with categorical and quantitative measures of flower intensity (intensity, -log_10_P = 16.28; brightness, -log_10_P = 19.2; chroma, -log_10_P = 18.08) (Figure 3b). We also infer the dark-associated allele at SNP1 has a partially dominant effect on flower color (brightness, dominance value (*d*/*a*) = 0.52; chroma, d/a = 0.3), while at SNP2 it has an additive effect (brightness, *d*/*a* = - 0.09; chroma, *d*/*a* = - 0.04).

Our full sequencing of this 35kb region combined with a rapid decay of LD around the most strongly associated variant supports SNP1 and SNP2 as candidate causal polymorphisms. Dark flower color was previously shown to be dominant to light flower color and is therefore expected to be controlled by a dominant mutation (Hopkins & Rausher, 2011; Billiard *et al*., 2021). Allelic variation at SNP1 meets this expectation with the dark associated allele being both exclusive to dark individuals and dominant in its predicted allelic effect. While variation at SNP2 alone cannot explain the evolution of dark flower color and its association with intensity may simply be an artifact of it being in LD with SNP1, we cannot yet rule out its involvement in affecting *PdMyb1* expression for dark color. Combined, these results indicate that a narrow non-coding region upstream of *PdMyb1* has a major effect on dark flower color, with a dominant point mutation in this region likely causing this effect. Notably, not all dark individuals carry the SNP1 dark-exclusive “G” allele. This observation suggests that multiple, possibly independently arising, mutations may cause convergent dark flower phenotype favored by reinforcement in sympatry.

### Dark associated mutations evolve new *cis*-regulatory motifs in the *PdMyb1* proximal promoter

Non-coding mutations can affect gene expression by modifying transcriptional regulation (Hill *et al*., 2021). We used *in silico* prediction to infer the positions of known *cis*-regulatory motifs within the 250bp upstream of *PdMyb1* from the *P. drummondii* reference genome. SNP1 and SNP2 are located between predicted CAAT-Box and TATA-Box elements of the *PdMyb1* core promoter (Figure 3d). This genomic region proximal to a gene’s translational start codon is known to have an important role in regulation and initiation of gene transcription (Hill *et al*., 2021; Schmitz *et al*., 2022).

Base pair changes at SNP1 and SNP2 are predicted to cause divergence in transcription factor binding sites inside the *PdMyb1* promoter (Figure 3d). The dark associated allele at SNP1 (“T”) gives rise to a putative antisense CAAT-Box-like binding motif ‘CCA**A**A” and optimizes the triple purine arm of a putative E2F TF binding motif ‘T**T**TGGCGG’ identified in dicots (Vandepoele *et al*., 2005). Neither of these motifs were predicted with the SNP1 light allele “G”. CAAT-boxes in antisense and “CCAAA” CCAAT-box derivatives can enhance transcription activation in plant promoters (Tiwari *et al*., 2010), and the presence\of an E2F TF binding site-like sequence in an R2R3-Myb promoter has been implicated in increased gene expression and flower color in roses (Li *et al*., 2022). The dark-associated allele at SNP2 (“A”) converts the binding site of a putative HD-Zip class I TF “CCAATGATTG” motif to that of a HD-Zip class II TF “CCAAT**A**ATTG” (Figure 3d), and this base change also modifies the DNA base following a CAAT-Box sequence “CCAAT**A**”. This base change is in positions known to modulate TF binding affinity and may alter recruitment of HD-Zip and CAAT-Box binding TFs (Bi *et al*., 1997; Ariel *et al*., 2007).

To determine the direction of evolution of the associated alleles at SNP1 and SNP2 we compared the sequences of the *PdMyb1* proximal promoter in dark sympatric and light allopatric *P. drummondii* individuals to representatives of the other annual *Phlox* species, *P. roemeriana* and *P. cuspidata*, and a perennial outgroup species, *P. pilosa ssp. pilosa* (Garner *et al*., 2022). Aside from the dark flowers observed in sympatric populations of *P. drummondii*, all four species display a light flower color. We find that the *PdMyb1* core promoter sequence is highly conserved across all species, with the three additional species homozygous for the *P. drummondii* light associated alleles at SNP1 (“G”) and SNP2 (“G”) (Figure 3e). This observation suggests the dark associated mutations at SNP1 (“T”) and SNP2 (“A”) arose within *P. drummondii*, leading to a gain of predicted *cis*-regulatory binding motif sequences in the dark *P. drummondii* promoter.

## Discussion

Reinforcement in *P. drummondii* is caused by the evolutionary transition from light to dark flower intensity (Hopkins & Rausher, 2011, 2012). We functionally demonstrate that the natural divergence in intensity is controlled by gene expression variation at an R2R3-Myb transcription factor, *PdMyb1*.

Evolution of novel *cis*-regulatory elements by a dominant point mutation in the *PdMyb1* promoter can explain the upregulation of *PdMyb1* expression and anthocyanin pigmentation in plants with dark flower color. Combined, our findings are the first functional demonstration of a causal gene contributing to speciation by reinforcement and the identification of the mutations and molecular mechanism responsible for its contribution to reproductive isolation in nature.

We have shown reinforcement can evolve quite simply, with a single mutation at a single gene explaining a large dominant phenotypic effect on flower color. This observation is consistent with theoretical models of reinforcement being able to successfully evolve despite gene flow when the genetic basis is simple and has a large phenotypic effect. This observation also supports the hypothesis that reinforcement may be more successful with a large effect dominant allele (Ortiz-Barrientos *et al*., 2004). Surprisingly, we demonstrate this adaptive leap can be explained by just a single base pair change at SNP1. While a second less strongly associated base change was observed at SNP2, it is also observed in light plants. This variant may have an epistatic effect on dark flower color conditional on the presence of the alleles at SNP1 or it may simply be associated by virtue of linkage. This discovery allows for future functional testing of alleles at both variants to identify their individual and combined contributions to dark flower color, their direct fitness effects in nature, and study of how selection acted on them to confer reinforcement. Additionally, mapping of the genetic basis of hybrid sterility between *P. drummondii* and *P. cuspidata* (i.e., the cost of hybridization) will be crucial to understanding how alleles at *PdMyb1* stay correlated with species-specific sterility alleles in the face of gene flow and recombination. Ultimately, more examples of mutations causing reinforcement traits are needed to determine if a simple mutational basis is common for successful evolution of this process.

The mutations at SNP1 and SNP2 alter the *cis*-regulatory and not the coding-sequence of *PdMyb1*. Our results are in line with *cis*-regulatory mutations of developmental regulatory genes causing adaptive morphological differences between species (Stern & Orgogozo, 2008; Martin & Orgogozo, 2013). Our findings also echo the trend observed in plants that these mutations are commonly found in gene promoters and alter pre-existing gene expression patterns (Della Pina *et al*., 2014). Our study suggests the genetic basis of reinforcement traits can evolve similarly to those for other adaptations maintaining species divergence, and our results motivate consideration of similar regulatory genetic changes as candidate causal variants in other reinforcement systems.

Growing evidence continues to link Subgroup 6 R2R3-Mybs to the divergence of floral anthocyanin concentration (intensity) and the evolution of premating isolation between young plant species pairs (Streisfeld & Rausher, 2010; Sobel & Streisfeld, 2013). Unlike other anthocyanin biosynthesis pathway regulating genes, Subgroup 6 R2R3-Mybs are often specialized in their expression profiles and developmental control. This specificity is believed to confer relatively minimal pleiotropic fitness effects, opening these genes up to being a substrate for evolution to diversify mating signals with limited cost. Linking *PdMyb1* function to dark flower color adds to the evidence of Subgroup 6 R2R3-Mybs as a hotspot for species diversification in plants and demonstrates reinforcement can evolve by similar pathways as premating divergence without selection against hybridization.

Known mutations causing variation in Subgroup 6 R2R3-Myb function have largely been *cis*-regulatory (Sheehan *et al*., 2016; Lin & Rausher, 2021), although see (Esfeld *et al*., 2018; Liang *et al*., 2022). These examples all involve a loss of function. However, we demonstrate evidence dark flower color may be a gain of function through the evolution of a novel transcription factor binding site upregulating *PdMyb1* expression and anthocyanin biosynthesis. Relative to losses, gains of novel traits are expected to be quite rare. It is possible the context of evolving reinforcement in *Phlox* specifically favored a gain of function mutation. R2R3-Myb loss of function mutations often cause a reduction or complete loss of flower color. Reduction in hybridization between *P. cuspidata* and *P. drummondii* in sympatry is due to the contrast between light and dark flower color inducing floral constancy in the foraging behavior of the *Battus philenor* butterfly (Hopkins & Rausher, 2012). Divergence to lighter or white *P. drummondii* flower color may not be a sufficiently large enough contrast from light flower colored *P. cuspidata* to induce this behavior. In this case, the success of reinforcement may have been contingent on how the direction and magnitude of flower color divergence is perceived by and manipulates a biological agent mediating reinforcing selection. Future study dissecting how the neurobiology of *B. philenor* interacts with flower color may aid in understanding how signal processing by a pollinator constrains or facilitates the phenotypic diversity and speciation of plants.

Our study demonstrates the power and promise of how innovations in functional genomics can elucidate the mutational basis of trait variation in non-model systems, connect it to the history and context of its evolution in nature, and resolve long standing questions on the origins of species. Continued development and applications of these approaches to isolating the genetic basis of other reproductive isolating traits, will yield a deepened understanding of species formation and the persistence of life on our planet.

## Materials and Methods

### Population sampling

We collected *P. drummondii* as seed from natural plant populations across the full species range in 2018, 2020, and 2021. This sampling included dark-red individuals from sympatry with *P. cuspidata*, light-blue individuals from allopatry with *P. cuspidata*, and natural genetically admixed individuals with recombinant flower colors from two independent zones where allopatry and sympatry meet (Supplemental Information Table S1). To preemptively control for ancestry, we grew only one individual from a known field maternal plant, preventing full sibs in our analyses. Seeds are submerged in a 500ppm Gibberellic acid solution for 48 hours at 4°C. Seeds are planted in Promix propogartion mix, placed in a dark 4*C for one week, and then allowed to germinate and grow under 23°C 16-hour day/ 18°C 8-hour night growing conditions.

### Phenotyping, Categorizing, and Quantifying Dimensions of Flower Color

All experimental individuals were phenotyped for flower color both quantitatively using spectral reflectance and qualitatively by eye into the 4 recombinant flower color phenotypes described in (Hopkins & Rausher, 2011). For each individual, raw spectral reflectance wavelengths were measured on the center of the petal lobe of twelve individual petals using a Flame-S-UV-VIS-ES spectrophotometer (Ocean Optics). The average of these reflectance spectra were decomposed into values along three axes of color space (i.e., hue, chroma, brightness) following the calculations in Smith, 2014. Under these formulas hue can be described as the position around the circumference of a circle in degrees. We adjusted the hue value in these formulas by 90° for all individuals, to avoid samples flanking the 0 ° /360° boundary.

We first applied an Anderson-Darling test for a normal distribution to the three axes of color variation to evaluate how many phenotypic clusters exist within the natural segregating variation of the admixed individuals. This test determined that the distributions of Hue (A = 18.868, p-value < 2.2e-16), Chroma (A = 6.2753, p-value = 1.72e-15), and Brightness (A = 2.9693, p-value = 1.697e-07) do not arise from a single normal gaussian distribution. We then applied unsupervised gaussian mixture modeling with Mclust v6.0.0 (Scrucca *et al*., 2016) to identify the optimal number of gaussian components (“clusters”) within these three dimensions of color space. Across 14 models varying in their geometric characteristics, Mclust identified the best fitting model to contain gaussian mixtures of four components (EEV, BIC= -391.4025) (Supplemental Information Table S2, Figure S1). The second (VEV, BIC= -399.7242) and third (EVV, BIC= -404.1648) best fitting models also support a mixture of four gaussian components under different model assumptions. Under the best fit model, the number of individuals classified as belonging to each component are N=30, N=46, N=24, N=34 (Supplemental Information Figure S2). We used an adjusted Rand index to compare the co-partitioning of individuals into groups between our unsupervised model and qualitative flower color classification. The scaled means and variances of the color axes of the four-component model indicate the presence of a light (high brightness, low chroma) and a dark (low brightness, high chroma) component in both the red and blue ranges of hue (Supplemental Information Figure S2, Table S3). We implemented a Chi squared test to compare the individual assignment to the model-inferred and qualitative-by-eye light-red, dark-red, light-blue, dark-blue clusters. The proportion of variation in hue, chroma, and brightness explained by classifying these gaussian components as light v. dark and red v. blue was determined by regression (Supplemental Information Figure S3). We used individual ANOVAs to determine if the inferred four-components significantly deviate in hue, brightness, and chroma from each other and from natural allopatric light-blue and sympatric dark-red plants (Supplemental Information Figure S4, Table S4a-b).

### Sequence-based isolation and characterization of *PdMyb1*

We used a BLASTn search of the partial coding sequences of the R2R3-Myb gene, *PdMyb1*, identified in Hopkins & Rausher, 2011 (GenBank ID: HQ127329.1-HQ127336.1) to query the location of the gene in the *Phlox drummondii* reference genome assembly V1.0 (unpublished data) (Camacho *et al*., 2009). We retrieved two hits with one nearly identical to the ∼440 bp partial coding sequences of *PdMyb1* (97-98% nucleotide similarity). *Ab initio* gene modeling of the 10kb sequence flanking the top BLAST hit with Fgenesh, with the *Primula vulgaris* genome as a reference, identified a 857 bp gene model for *PdMyb1* (Solovyev *et al*., 2006).

We aligned the 268 amino acid sequence of this gene model to those of known anthocyanin-regulating R2R3-Myb genes from Arabidopsis, Antirrhinum, Grape, and Petunia (GenBank ID: NP_176813.1, AKB94073.1, ABB83827.1, BAD18977.1, ADW94950.1) (Supplemental Information Figure S5). With this alignment, we identified and annotated well-known conserved motifs of anthocyanin-regulating dicot R2R3-Myb transcription factors; the bhLH interacting motif [DE]Lx_2_[RK]x_2_Lx_6_Lx_3_R, a conserved dicot anthocyanin-promoting R3 motif [A/S/G]NDV, and the C-terminal R2R3-Myb SG6-defining motif [R/K]Px[P/A/R]x2[F/Y/L/R] (Stracke *et al*., 2001; Zimmermann *et al*., 2004; Lin-Wang *et al*., 2010; Hichri *et al*., 2011; Berardi *et al*., 2021).

### Virus induced gene silencing of *PdMyb1*

We acquired pTRV1 and pTRV2-MCS constructs from the Arabidopsis Biological Resource Center (ABRC accessions CD3-1039, CD3-1040, respectively). Unique sequences of the coding regions of the *Phlox drummondii PdMyb1* (307bp) and phytoene desaturase (*PdPDS*) (310bp) genes flanked with BamHI and EcoRI digestion sites were synthesized using Twist Biosciences (South San Francisco, California) (Supplemental Information Table S7). These synthesized fragments were independently cloned into the pTRV2-MCS plasmid using the BamHI and EcoRI digestion sites.

Our pTRV2-PdMyb1, pTRV2-PdPDS, and empty pTRV1 vectors were independently transformed into *Agrobacterium tumefasciens* (strain GV3101) and allowed to grow at 28*C overnight on YEB agar plates with kanamycin, gentamycin, and rifampicin antibiotic resistance selection. After PCR confirmation for the pTRV vectors, single colonies were used to inoculate 3mL overnight YEB cultures with kanamycin and gentamycin resistance selection. Cultures were grown overnight at 28*C and then used to initiate 50mL volume overnight cultures the following day. Cells were harvested by pelleting via centrifugation, resuspended in freshly made infiltration buffer (10mM MES, 200uM acetosyringone, 10mM MgCl+2), and normalized to OD_600_=10. pTRV2-PdMyb1 and pTRV2-PdPDS resuspended cells were separately mixed in a 1:1 volume with pTRV1 resuspended cells. Cell mixtures were incubated at room temperature for 2.5 hours before transfection.

*P. drummondii* seeds were germinated and grown under controlled conditions in a growth chamber. Once the plants began to branch, they were transfected with either our pTRV2-PdMyb1 + pTRV1 or pTRV2-PdPDS + pTRV1 inoculums by both applying the mixture to cut apical meristems and by injection into axillary buds with a 28-gauge needle. We transfected a total of 41 individuals with pTRV2-PdMyb1 + pTRV1 and 30 individuals with pTRV2-PdPDS1 + pTRV1. After transfection, all plants were placed into a growth chamber under 23°C 16-hour day/ 18°C 8-hour night growing conditions, with all plants under a plastic propagation domed for the first 24 hours.

Symptoms of virus-induced gene silencing appeared between 3-4 weeks after transfection. Single branches on each transfected plant were identified by eye to be silenced based on photobleaching of the vegetative tissue (pTRV2-PdPDS) or color change of the floral tissue (pTRV2-PdMyb1). For each branch, flower color was phenotyped by spectral reflectance as described above. For both branch types, RNA was extracted from a pool of three stage six buds and converted to cDNA as described above. We then used quantitative real time PCR to quantify expression of *PdMyb1* and the housekeeping gene *PdEF1a* in pTRV2-PdMyb1 in silenced (N=20) and unsilenced (N=20) branches, pTRV2-PdPDS silenced branches (N=10), and no transfection control wildtype dark (N=13) and light (N=13) plants. cDNA from a single sample was run across all plates to normalize for batch effects. Significance differences between groups was determined using ANOVAs with post hoc Tukey test (Figure 2b-d, Supplemental Information Table S5a-d).

### Targeted enrichment of PdMyb1 genomic region, long-read sequencing, and local variant identification

We isolated the genomic window around the *PdMyb1* gene using the X-drop™ method for target enrichment of high molecular weight (HMW) long fragment DNA (Madsen *et al*., 2020). Genome-wide high molecular weight nuclear DNA was extracted from Phlox meristem tissue using a modified nuclei isolation and chloroform extraction method. We targeted *PdMyb1* containing HMW DNA fragments with primers specific to the C-terminus of the gene’s third exon (d100227_1_3_F: 5’ACATAGCGGGAGTCATTGGG3’, d100227_1_3_R: 5’ACCTCTATCCCTGGTACCGT3’) and amplified isolated HMW DNA positive for *PdMyb1* was amplified using multiple displacement amplification (MDA). Oxford Nanopore Technologies (ONT) sequencing libraries were constructed and barcoded by individual and sequenced using an ONT PromethION with R9.4.1 flowcells. The Xdrop enrichment and Nanopore sequencing was performed by Samplix ApS services (Herlev, Denmark).

Raw long-read sequence data was basecalled and demultiplexed with GUPPY v.5.0.11 HAC model (Oxford Nanopore Technologies). 10bp from the head and tail of each read was clipped, and reads were filtered for an average base quality ≥ 10 and length ≥ 500bp using Nanofilt (De Coster *et al*., 2018). Trimmed and filtered long reads were error corrected using Canu (version 1.8) with settings - corOutCoverage =1000 -corMinCoverage = 0 (Koren *et al*., 2017), and reads with chimeric structure resulting from MDA were split using SACRA with settings sl=1000, pc=10 (Kiguchi *et al*., 2021), per recommendation by Samplix ApS.

To identify simple variants (SNPs and small indels), our processed reads were mapped to the *P. drummondii* reference genome assembly V1.0 (unpublished data) using minimap2 v2.1 (Li, 2018) with settings -z 600,200 -x map-ont. Variants within a 6Mb window flanking the R2R3-Myb1 gene (i.e. chrlg6:19000000-26000000) were called using Clair3 with the “enable long indel” parameter (Zheng *et al*., 2021).

### Region-wide genotype-phenotype association analyses

A joint cohort-level gVCF was constructed using GLNexus (Lin *et al*., 2018; Yun *et al*., 2021), and variants were filtered using bcftools (Danecek *et al*., 2021) to be 1) polymorphic, 2) Depth >10X, 3) GQ >10, 4) MAF >0.025, and 5) have <90% missingness. After filtering, 2,533 variants were left for association analyses. Genotype-phenotype associations were performed using a univariate linear mixed model in GEMMA (version 0.98.3) (Zhou & Stephens, 2012). Population structure and relatedness among individuals were accounted for in our association analyses using a relatedness matrix generated by GEMMA from our region-wide variants. Significant associations with categorical and quantitative measures of (light versus dark) and quantitative brightness and chroma phenotypes were evaluated using the P value of a two-sided Wald test, with a Bonferroni correction for multiple testing (cutoff = −log_10_ (0.05/number of variants). We used the *d*/*a* statistic to infer the dominance relationship of alleles at each significantly associated SNP (Miller *et al*., 2014). The allelic effect was categorized based on the *d*/*a* ratio falling into one of the following categories: < −1.25 for underdominant, −1.25 to −0.75 for recessive, −0.75 to −0.25 for partially recessive, −0.25 to 0.25 for additive, 0.25 to 0.75 for partially dominant, 0.75 to 1.25 for dominant, and >1.25 for overdominant.

### Annotation of *cis*-regulatory elements

*Cis*-regulatory element motifs within 250bp upstream of *PdMyb1* were inferred using PlantPAN 3.0 (Chow *et al*., 2019) and PlantCARE (Lescot *et al*., 2002) with the *P. drummondii* reference genome sequence.

## Supporting information

Supplemental Information for Garner Mutation

