## Supplemental Information for Garner Mutation for "A *cis*-regulatory point mutation at a R2R3-Myb transcription factor contributes to speciation by reinforcement in *Phlox drummondii*"

**Supplemental information Table S3.1.** Populations from which *P. drummondii* seeds were collected.

| Location | Population | Coordinates |
|---|---|---|
| Northern Admixed | 2020_01 | (30.06752, -97.21527) |
| Northern Admixed | 2020_02 | (30.13561, -97.37201) |
| Northern Admixed | 2020_03/ 2020_08 | (30.0665, -97.3526) |
| Northern Admixed | 2020_07 | (30.05097, -97.23811) |
| Northern Admixed | 2020_09 | (30.16084, -97.3907) |
| Northern Admixed | 764 | (30.10343, -97.22129) |
| Northern Admixed | 770 | (30.20639, -97.34583) |
| Southern Admixed | 671 | (29.4161, -97.70415) |
| Southern Admixed | 672 | (29.41633, -97.71911) |
| Southern Admixed | 673 | (29.39921, -97.74658) |
| Southern Admixed | 674 | (29.40755, -97.75621) |
| Southern Admixed | 675 | (29.4024, -97.77451) |
| Southern Admixed | 678/759 | (29.28641, -97.6241) |
| Allopatry | 702d | (30.90258, -99.28651) |
| Allopatry | 703b | (30.7888, -99.31741) |
| Allopatry | 704b | (30.75833, -99.0579) |
| Allopatry | 706b | (30.63676, -98.4487) |
| Allopatry | 777 | (29.23307, -98.64285) |
| Allopatry | 778 | (28.96803, -98.47288) |
| Allopatry | 785 | (30.27233, -98.84158) |
| Sympatry | 680a | (29.73633, -97.41014) |
| Sympatry | 656a | (30.06134, -96.92056) |
| Sympatry | 667 | (29.52881, -96.44123) |
| Sympatry | 802 | (29.71848, -96.8887) |
| Sympatry | 831 | (29.86051, -97.59049) |

**Supplemental Table S3.2.** Bayesian Information Criterion (BIC) values for Mclust finite gaussian mixture models fit to natural *P. drummondii* flower color variation. Mclust fit 14 different geometric mixture models with 1 to 9 underlying gaussian components. The five most optimal mixture models (lowest BIC values) are indicated in bold.

| Mixture model | Number of gaussian components | | | | | | | | |
|---|---|---|---|---|---|---|---|---|---|
| | 1 | 2 | 3 | 4 | 5 | 6 | 7 | 8 | 9 |
| EII | -1157.41 | -923.36 | -835.30 | -607.13 | -561.62 | -517.28 | -526.62 | -504.94 | -475.35 |
| VII | -1157.41 | -843.58 | -774.64 | -611.91 | -558.42 | -525.51 | -528.89 | -522.59 | -506.72 |
| EEI | -1167.20 | -890.47 | -823.46 | -573.01 | -550.07 | -502.98 | -512.29 | -500.71 | -466.92 |
| VEI | -1167.20 | -843.56 | -740.50 | -581.04 | -543.22 | -512.72 | -522.85 | -510.86 | -495.02 |
| EVI | -1167.20 | -883.95 | -817.83 | -559.07 | -552.98 | -506.16 | -522.96 | -530.72 | -500.81 |
| VVI | -1167.20 | -802.42 | -739.35 | -564.20 | -544.93 | -519.16 | -528.93 | -540.54 | -533.29 |
| EEE | -735.62 | -666.66 | -482.07 | -426.96 | -430.19 | -426.65 | -434.22 | -437.44 | -429.89 |
| VEE | -735.62 | -652.03 | -484.91 | -439.34 | -443.25 | -446.54 | -460.47 | -464.63 | -462.41 |
| EVE | -735.62 | -555.65 | -463.92 | -423.73 | -441.21 | -443.18 | -457.84 | -470.96 | -483.36 |
| VVE | -735.62 | -571.60 | -463.59 | -436.51 | -450.65 | -463.35 | -481.23 | -501.12 | -514.94 |
| EEV | -735.62 | -644.93 | -442.55 | **-391.40** | **-406.61** | -425.10 | -445.86 | -469.77 | -490.14 |
| VEV | -735.62 | -595.38 | -432.91 | **-399.72** | -416.56 | -439.84 | -468.20 | -493.12 | -511.10 |
| EVV | -735.62 | -492.77 | -447.96 | **-404.16** | -427.52 | -451.58 | -497.24 | -522.81 | -550.04 |
| VVV | -735.62 | -575.14 | -433.45 | **-415.28** | -440.22 | -472.93 | -512.81 | -551.55 | -576.01 |

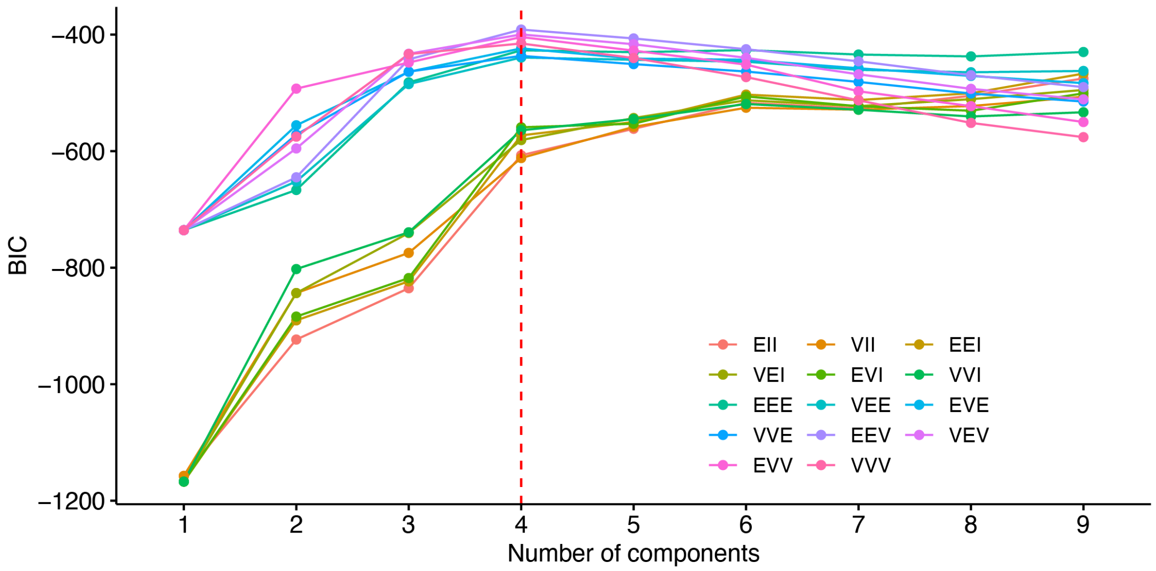


**Supplemental Information Figure S3.1.** Bayesian Information Criterion (BIC) values for Mclust finite gaussian mixture models fit to natural *P. drummondii* flower color variation. Optimal model is EEV with 4 underlying gaussian components.

**Supplemental Information Figure S3.2**. Density and classification of individuals under Mclust best fit mixture of four-component model relative to scaled values of hue, chroma, and brightness. Individual classification to each component is indicated as either pink circles (component 1), blue squares (component 2), blue triangles (component 3), or red crosses (component 4).

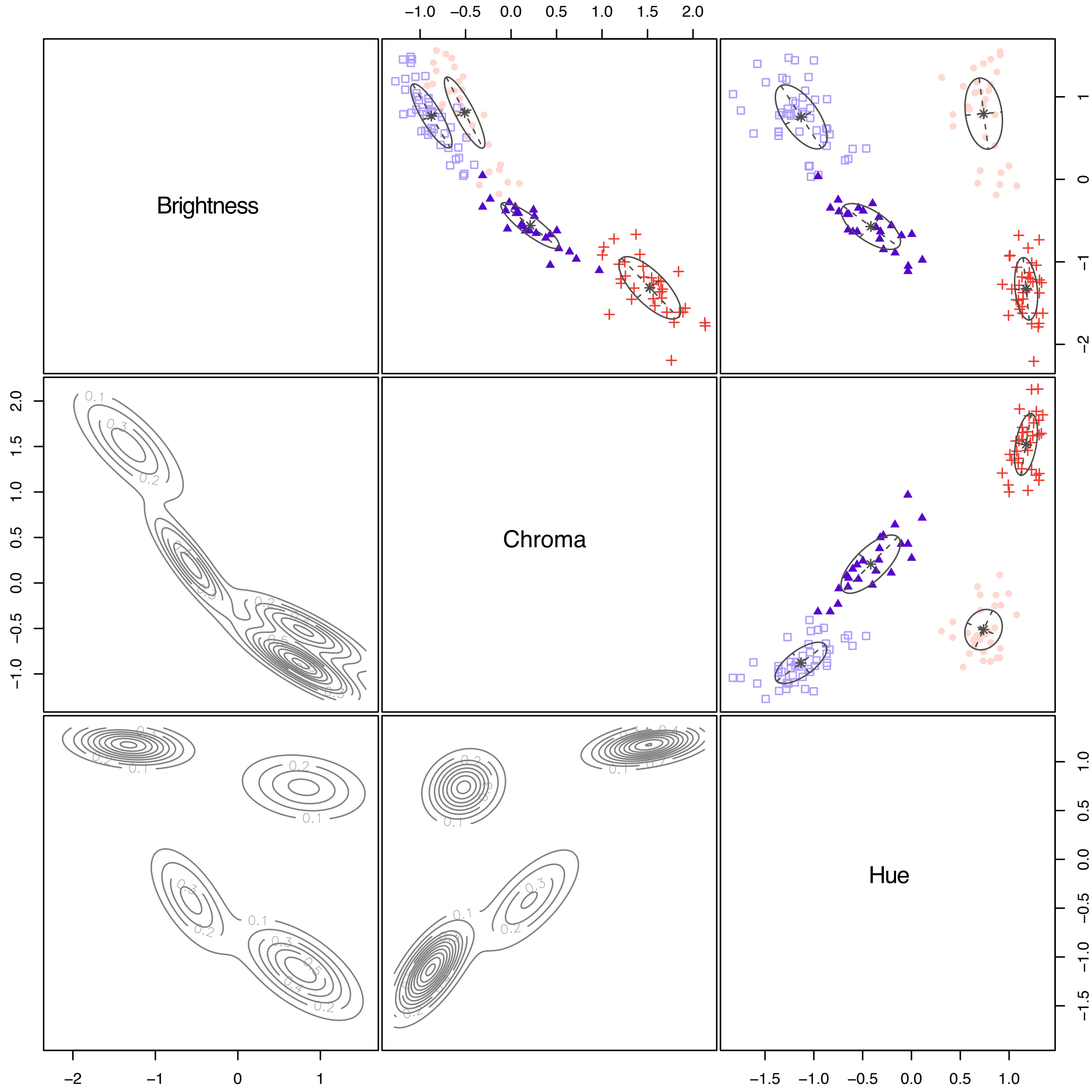

**Supplemental Table S3.3.** Values of hue, chroma, and brightness of the four model-inferred gaussian components and their inferred categorical phenotype.

| Component | Brightness | | Hue | | Chroma | | Inferred Categorical Phenotype |
|---|---|---|---|---|---|---|---|
| | Mean | Standard Error | Mean | Standard Error | Mean | Standard Error | |
| 1 | 14354.400 | 273.344 | 90.695 | 0.379 | 0.359 | 0.008 | Light Red |
| 2 | 14282.93 | 153.6818 | 68.408 | 0.504 | 0.297 | 0.005 | Light Blue |
| 3 | 10542.2 | 172.771 | 76.690 | 0.737 | 0.475 | 0.011 | Dark Blue |
| 4 | 8337.381 | 160.031 | 95.868 | 0.226 | 0.695 | 0.008 | Dark Red |

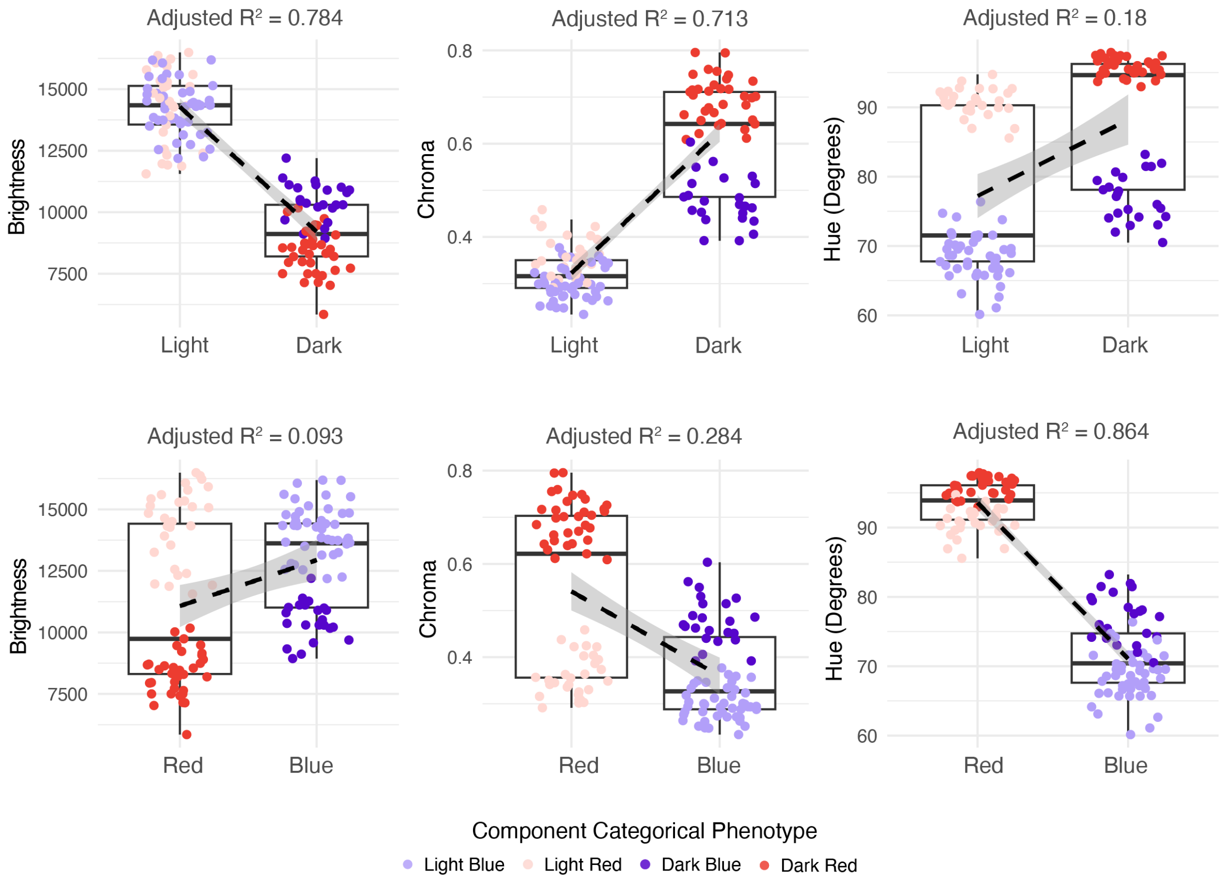


**Supplemental Information Figure S3.3**. Linear regressions of spectral reflectance axes by categorical light v. dark or red v. blue flower color for the four model-inferred gaussian components. All regressions are significant, P < 0.001.

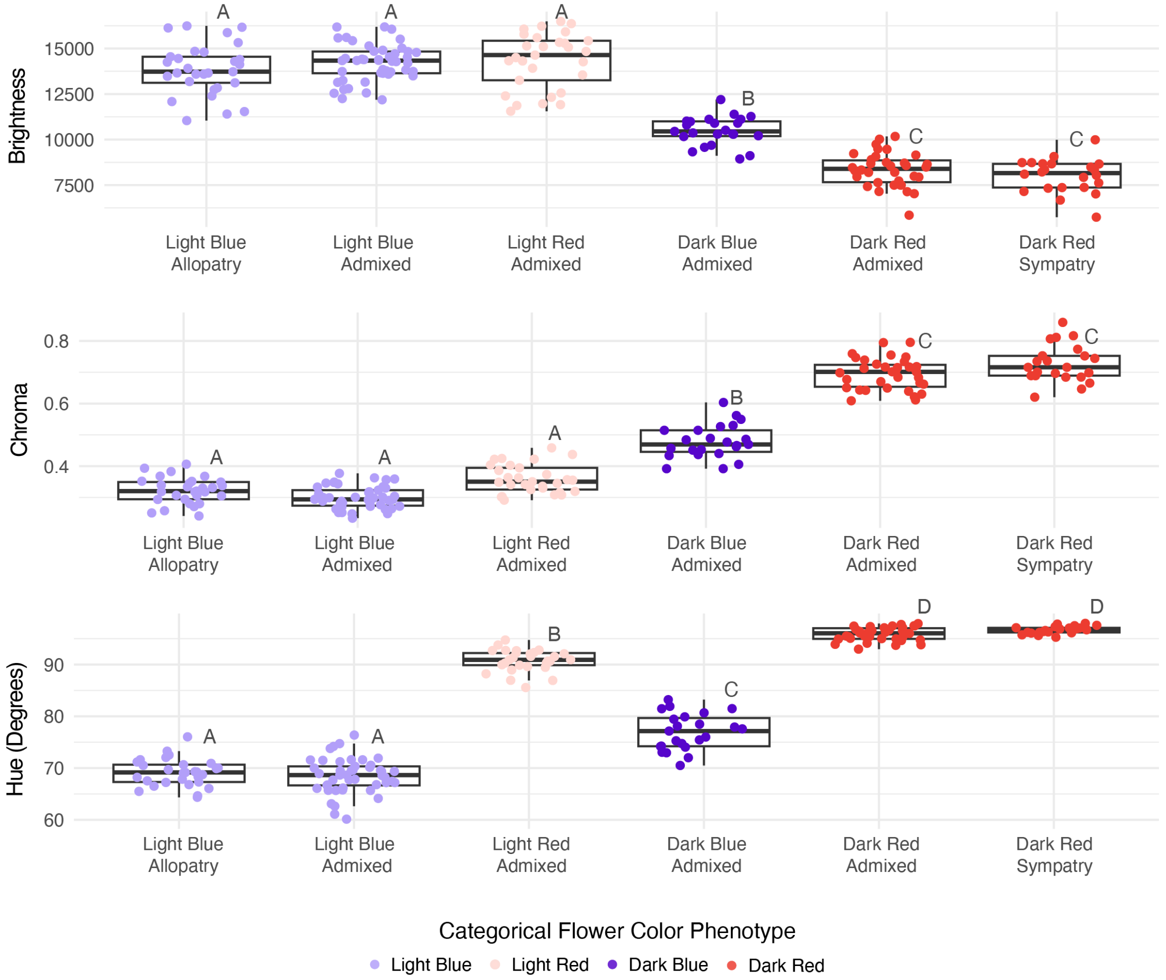


**Supplemental Information Figure S3.4**. Four-components significantly deviate in hue, brightness, and chroma from each other, with dark-red and light-blue admixed individuals resembling the natural spectral reflectance of allopatric light-blue and sympatric dark-red plants. Letters denote significant differences between groups, as determined with ANOVAs with the threshold of $P < 0.001$ (Supplemental Information Table S3.4a-b).

**Supplemental Information Table S3.4a.** ANOVA results for comparison between Brightness, Chroma, and Hue between the four model-inferred gaussian components and allopatric and sympatric plant groups.

| Dependent Variable | Hue | | | Brightness | | | Chroma | | |
|---|---|---|---|---|---|---|---|---|---|
| | df | F | P | df | F | P | df | F | P |
| Group | 5 | 816.900 | <2e-16 | 5 | 214.300 | <2e-16 | 5 | 524.900 | <2e-16 |

**Supplemental Information Table S3.4b.** Post hoc Tukey tests of ANOVA comparison of brightness, chroma, and hue between the admixed four model-inferred gaussian components (LB, LR, DB, DR) and allopatric light-blue (Allo-LB) and sympatric dark-red (Sym-DR) plant groups.

| Independent Variable | Treatment (I) | Treatment (J) | Mean Difference (I-J) | Lower | Upper | Adjusted P |
|---|---|---|---|---|---|---|
| Brightness | LB | DB | 3740.729 | 2927.591 | 4553.868 | <0.001 |
| | | DR | 5945.548 | 5204.819 | 6686.278 | <0.001 |
| | | Sym-DR | 6280.786 | 5445.229 | 7116.343 | <0.001 |
| | LR | DB | 3812.198 | 2929.438 | 4694.958 | <0.001 |
| | | Sym-DR | 6352.254 | 5448.802 | 7255.707 | <0.001 |
| | | DR | 6017.017 | 5200.469 | 6833.565 | <0.001 |
| | | LB | 71.469 | -696.875 | 839.812 | 1.000 |
| | | Allo-LB | 512.054 | -336.849 | 1360.957 | 0.509 |
| | Allo-LB | DB | 3300.144 | 2410.493 | 4189.795 | <0.001 |
| | | DR | 5504.962 | 4680.969 | 6328.956 | <0.001 |
| | | LB | -440.586 | -1216.837 | 335.666 | 0.577 |
| | | Sym-DR | 5840.200 | 4930.013 | 6750.387 | <0.001 |
| | DR | DB | -2204.819 | -3063.651 | -1345.986 | <0.001 |
| | Sym-DR | DB | 2540.056 | -3481.899 | -1598.213 | <0.001 |
| | | DR | -335.238 | -1215.325 | 544.850 | 0.882 |
| Chroma | LB | DB | -0.178 | -0.211 | -0.145 | <0.001 |
| | | DR | -0.398 | -0.428 | -0.368 | <0.001 |
| | | Sym-DR | -0.431 | -0.465 | -0.397 | <0.001 |
| | LR | DB | -0.116 | -0.152 | -0.080 | <0.001 |
| | | Sym-DR | -0.369 | -0.405 | -0.332 | <0.001 |
| | | DR | -0.336 | -0.369 | -0.302 | <0.001 |
| | | LB | 0.062 | 0.031 | 0.094 | <0.001 |
| | | Allo-LB | 0.037 | 0.002 | 0.072 | 0.028 |
| | Allo-LB | DB | -0.153 | -0.189 | -0.117 | <0.001 |
| | | DR | -0.373 | -0.406 | -0.339 | <0.001 |
| | | LB | 0.025 | -0.006 | 0.057 | 0.195 |
| | | Sym-DR | 0.025 | -0.006 | 0.057 | 0.195 |

**Supplemental Information Table S3.4b Continued.** Post hoc Tukey tests of ANOVA comparison of Brightness, Chroma, and Hue between the admixed four model-inferred gaussian components (LB, LR, DB, DR) and allopatric light-blue (Allo-LB) and sympatric dark-red (Sym-DR) plant groups.

| Independent Variable | Treatment (I) | Treatment (J) | Mean Difference (I-J) | Lower | Upper | Adjusted P |
|---|---|---|---|---|---|---|
| | Sym-DR | DB | 0.253 | 0.214 | 0.291 | <0.001 |
| | | DR | 0.033 | -0.003 | 0.069 | 0.092 |
| Hue | LB | DB | -8.282 | -10.151 | -6.413 | <0.001 |
| | | DR | -27.460 | -29.163 | -25.757 | <0.001 |
| | | Sym-DR | 28.343 | -30.264 | -26.422 | <0.001 |
| | LR | DB | 14.005 | 11.976 | 16.034 | <0.001 |
| | | Sym-DR | -6.056 | -8.133 | -3.979 | <0.001 |
| | | DR | -5.173 | -7.050 | -3.296 | <0.001 |
| | | LB | 22.287 | 20.521 | 24.053 | <0.001 |
| | | Allo-LB | 21.585 | 19.634 | 23.536 | <0.001 |
| | Allo-LB | DB | -7.580 | -9.625 | -5.535 | <0.001 |
| | | DR | -26.758 | -28.652 | -24.864 | <0.001 |
| | | LB | 0.702 | -1.082 | 2.486 | <0.001 |
| | | Sym-DR | -27.641 | -29.733 | -25.549 | <0.001 |
| | DR | DB | 19.178 | 17.204 | 21.152 | <0.001 |
| | Sym-DR | DB | 20.061 | 17.896 | 22.226 | <0.001 |
| | | DR | 0.883 | -1.140 | 2.906 | 0.808 |

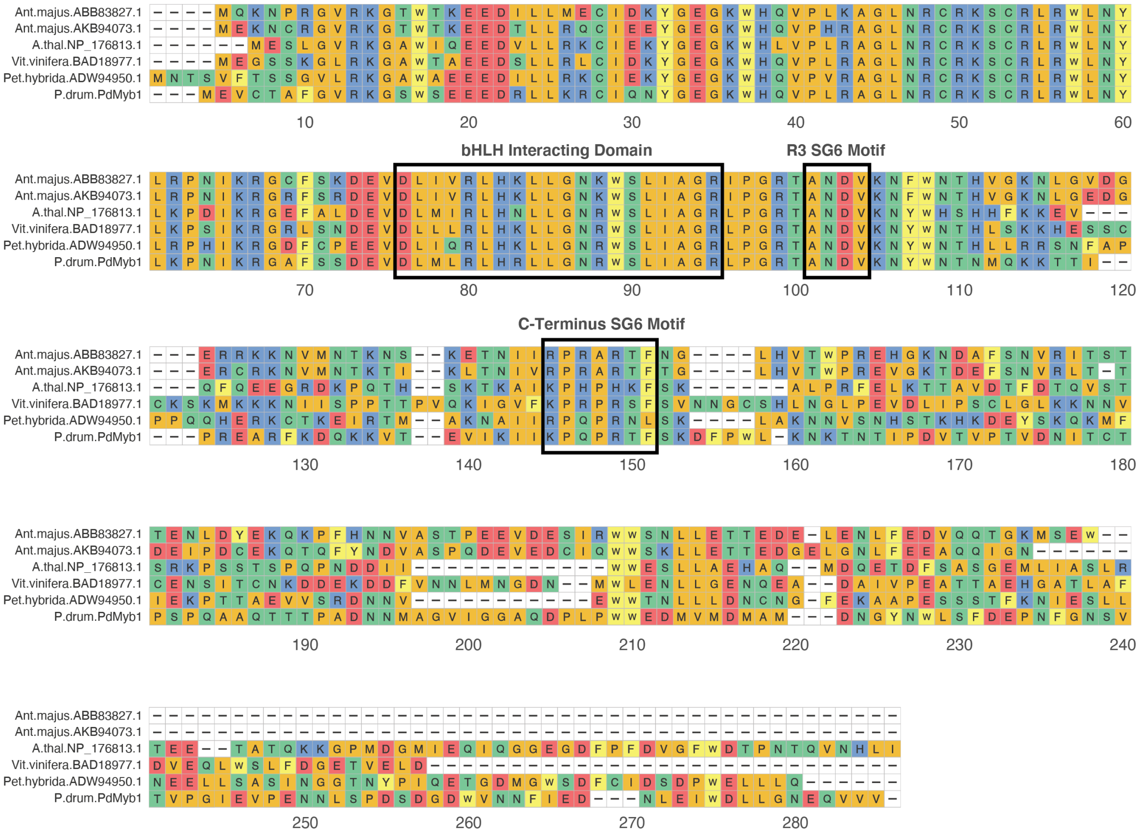


**Supplemental Information Figure S3.5**. Amino acid alignment of *PdMyb1* and known anthocyanin-regulating subgroup 6 (SG6) R23R3-Myb transcription factors. Boxes indicate three highly conserved SG6 R2R3-Myb amino acid sequences; a bHLH interacting domain, R3 SG6 Motif, and C-Terminus SG6 motif are indicated.

**Supplemental Information Table S3.5.** Results for *PdMyb1* expression, brightness, and chroma in independent treatment groups ANOVAs. ANOVAs were repeated so paired VIGS treatments *PdMyb1* Silenced and *PdMyb1* Unsilenced can be considered separately.

| VIGS Treatment | Dependent Variable | *PdMyb1* expression | | | Brightness | | | Chroma | | |
|---|---|---|---|---|---|---|---|---|---|---|
| | | df | F | P | df | F | P | df | F | P |
| PdMyb1 Silenced | Treatment | 3 | 31.770 | 8.3E-12 | 3 | 121.200 | <2E-16 | 3 | 61.180 | <2E-16 |
| PdMyb1 Unsilenced | Treatment | 3 | 11.470 | 6.97E-06 | 3 | 204.900 | <2E-16 | 3 | 57.270 | <2E-16 |

**Supplemental Information Table S3.5b.** Post hoc Tukey test for independent VIGS treatment groups ANOVAs including *PdMyb1* Silenced

| Independent Variable | Treatment (I) | Treatment (J) | Mean Difference (I-J) | Lower | Upper | Adjusted P |
|---|---|---|---|---|---|---|
| *PdMyb1* | Dark Untreated | *PdMyb1* Silenced | 0.718 | 0.437 | 1.000 | <0.001 |
| Expression | Light Untreated | *PdMyb1* Silenced | 0.125 | -0.157 | 0.406 | 0.645 |
| | | Dark Untreated | -0.594 | -0.904 | -0.284 | <0.001 |
| | *PdPDS* Silenced | *PdMyb1* Silenced | 0.948 | 0.642 | 1.254 | <0.001 |
| | | Dark Untreated | 0.230 | -0.102 | 0.562 | 0.268 |
| | | Light Untreated | 0.824 | 0.492 | 1.156 | <0.001 |
| Brightness | Dark Untreated | *PdMyb1* Silenced | -2887.759 | -3974.440 | -1801.079 | <0.001 |
| | Light Untreated | *PdMyb1* Silenced | 4003.077 | 2916.396 | 5089.757 | <0.001 |
| | | Dark Untreated | 6890.836 | 5694.440 | 8087.232 | <0.001 |
| | *PdPDS* Silenced | *PdMyb1* Silenced | -4206.828 | -5388.174 | -3025.483 | <0.001 |
| | | Dark Untreated | -1319.069 | -2602.060 | -36.078 | 0.042 |
| | | Light Untreated | -8209.905 | -9492.896 | -6926.914 | <0.001 |
| Chroma | Dark Untreated | *PdMyb1* Silenced | 0.197 | 0.120 | 0.275 | <0.001 |
| | Light Untreated | *PdMyb1* Silenced | -0.154 | -0.231 | -0.076 | <0.001 |
| | | Dark Untreated | -0.351 | -0.436 | -0.266 | <0.001 |
| | *PdPDS* Silenced | *PdMyb1* Silenced | 0.238 | 0.154 | 0.322 | <0.001 |
| | | Dark Untreated | 0.041 | -0.050 | 0.132 | 0.633 |
| | | Light Untreated | 0.392 | 0.301 | 0.483 | <0.001 |

**Supplemental Information Table S3.5c.** Post hoc Tukey test for independent VIGS treatment groups ANOVAs including *PdMyb1* Unsilenced

| Independent Variable | Treatment (I) | Treatment (J) | Mean Difference (I-J) | Lower | Upper | Adjusted P |
|---|---|---|---|---|---|---|
| *PdMyb1* Expression | *PdMyb1* Unsilenced | Dark Untreated | 0.018 | -0.330 | 0.366 | 0.999 |
| | | Light Untreated | 0.612 | 0.264 | 0.960 | <0.001 |
| | Light Untreated | Dark Untreated | -0.594 | -0.977 | -0.211 | <0.001 |
| | *PdPDS* Silenced | *PdMyb1* Unsilenced | 0.212 | -0.167 | 0.590 | 0.454 |
| | | Dark Untreated | 0.230 | -0.181 | 0.641 | 0.453 |
| | | Light Untreated | 0.824 | 0.413 | 1.235 | <0.001 |
| Brightness | *PdMyb1* Unsilenced | Dark Untreated | -229.378 | -1118.604 | 659.848 | 0.902 |
| | | Light Untreated | -7120.214 | -8009.440 | -6230.988 | <0.001 |
| | Light Untreated | Dark Untreated | 6890.836 | 5911.830 | 7869.842 | <0.001 |
| | *PdPDS* Silenced | *PdMyb1* Unsilenced | -1089.691 | -2056.382 | -123.001 | 0.021 |
| | | Dark Untreated | -1319.069 | -2368.936 | -269.203 | 0.008 |
| | | Light Untreated | -8209.905 | -9259.772 | -7160.039 | <0.001 |
| Chroma | *PdMyb1* Unsilenced | Dark Untreated | -0.002 | -0.085 | 0.080 | 1.000 |
| | | Light Untreated | 0.348 | 0.266 | 0.431 | <0.001 |
| | Light Untreated | Dark Untreated | -0.351 | -0.441 | -0.260 | <0.001 |
| | *PdPDS* Silenced | *PdMyb1* Unsilenced | 0.043 | -0.046 | 0.133 | 0.575 |
| | | Dark Untreated | 0.041 | -0.056 | 0.138 | 0.679 |
| | | Light Untreated | 0.392 | 0.295 | 0.489 | <0.001 |

**Supplemental Information Table S3.6.** Results for paired T-tests between VIGS Treatments *PdMyb1* Silenced and *PdMyb1* Unsilenced for *PdMyb1* expression, Brightness, and Chroma.

| *PdMyb1* Silenced - *PdMyb1* Unsilenced Paired Comparison | Mean | 95% confidence interval | | df | T-test Value | P |
|---|---|---|---|---|---|---|
| | | Lower | Upper | | | |
| *PdMyb1* expression | -0.737 | -0.900 | -0.573 | 19 | -9.455 | 1.29E-08 |
| Brightness | 3117.137 | 2416.924 | 3817.350 | 19 | 9.318 | 1.623E-08 |
| Chroma | -0.195 | -0.228 | -0.162 | 19 | -12.498 | 1.30E-10 |

**Supplemental Information Table S3.7.** Targeted coding sequences for virus-induced gene silencing in *P. drummondii.*

| Gene Target | Sequence 5’-3’ |
|---|---|
| *PdPDS* | ATGAGTACGAATTCTTGGAGTTCCAGTTATAAATATCCACATATGGTTTGACAG<br>AAAACTGAAGAACACATACGACCACCTTCTTTTCAGCAGAAGTCCCCTTCTCAG<br>TGTATATGCTGACATGTCTCTCACATGCAAGGAATATTATGACCCAAACAAATC<br>TATGCTGGAATTGGTGTTTGCACCTGCAGAGGAATGGATCAACCGTAGTGACG<br>AAGAAATTATTGATGCTACAATGATGGAACTTTCAAAACTCTTTCCTGATGAAAT<br>TTCTGCAGATCAAAGCAAAGCGAAAATATTGAAGTACAAAGTCGGGATCCCAG<br>CCTGA |
| *PdMyb1* | ATGAGTACGAATTCCCACCACCACAAGCAGCCCAAACAGCAACACCAGCCGA<br>CAATAACATAGCGGGAGTCATTGGGGTTGCTCAAGACCCATTACCATGGTGGG<br>AAGACATGGTGATGGACATGGGAATGGATAATGGCAACAATTGGTTGTCCTTT<br>GATGAACCCAATTTCGGGAATAGTGTAACGGTACCAGGGATAGAGGTACCTGA<br>GAATAACTTGTCCCCGGATAGCGATGGAGATTGGGTCAATAATTTCATTGAGG<br>ATAACTTGGAGATATGGGATCTTCTAGGTAACGAACAAGTTGTGGTCTAAGGAT<br>CCCAGCCTGA |

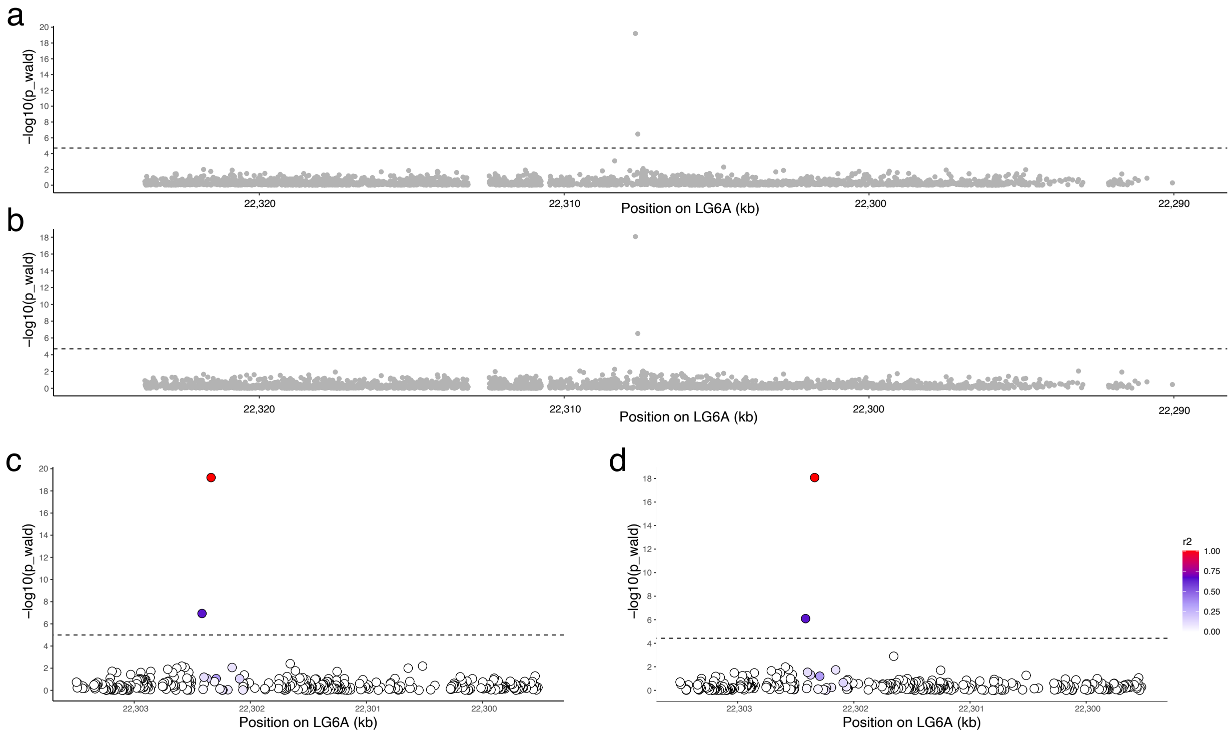


**Supplemental Information Figure S3.6**. Genetic associations with natural variation in *P. drummondii* quantitative flower color brightness **(a,c)** and chroma **(b,d)** identify only two associated variants the 35kb window flanking *PdMyb1*. The dashed line represents the Bonferroni corrected significance threshold ((-log$_{10}$(P) $\simeq$4.7). Circles represent genetic variants. Colors in lower c and d represent the degree of linkage disequilibrium between each marker and the highest associated variant, SNP1.
